# Modelling of brain dynamics reveals reduced switching between brain states in insomnia disorder - a resting state fMRI study

**DOI:** 10.1101/2024.11.27.625644

**Authors:** Kira Vibe Jespersen, Angus Stevner, Morten Kringelbach, Eus Van Someren, Diego Vidaurre, Peter Vuust

## Abstract

Insomnia disorder is the most common sleep disorder, and neuroimaging research indicates that it is related to dysfunction in large-scale brain networks. Recently developed methods have enabled the investigation of the dynamic aspects of brain activity varying over time. In the present study, we used a novel data-driven approach to evaluate time-varying brain activity in adults with insomnia disorder compared to matched controls with no sleep problems. We acquired ten minutes resting state functional magnetic resonance images and T1-weighed images in all participants. We used Hidden Markov modelling for a data-driven definition of dynamic changes in whole-brain activity. The results showed that insomnia disorder is characterised by reduced switching rates between brain states. In line with the reduced switching, the HMM analyses suggested reduced prevalence of two whole-brain states – the default mode network and a fronto-parietal network – and an increase in just one brain state – a global activation state – in insomnia patients compared to controls. The findings suggest that insomnia disorder is characterised by less flexible transitions between brain states at wakeful rest, and thus highlight the importance of evaluating the spatiotemporal dynamics of brain activity to advance the understanding of the neural underpinnings of insomnia disorder.

## Introduction

Insomnia is the most common sleep disorder characterised by persistent difficulties initiating or maintaining sleep accompanied by impaired daytime function (Riemann et al., 2017). It is the second-most common mental disorder in Europe (Wittchen et al., 2011) and prevalence estimates vary between 6% and 20% (Ohayon, 2002; Riemann et al., 2017) with higher prevalence in women and older adults. The prevailing model of the underlying mechanism of insomnia is the hyper-arousal theory, stating that insomnia is the result of a 24-hour increased arousal as reflected in both psychological and neurophysiological measures.

An increasing number of studies have examined the neural correlates of insomnia, but the results vary considerably (Aquino et al., 2024; Tahmasian et al., 2018). Magnetic resonance imaging studies suggest insomnia-related dysfunction in large-scale brain networks (Jespersen et al., 2020; Kay & Buysse, 2017; Khazaie et al., 2017; Van Someren, 2021). Several studies have used resting-state functional magnetic resonance imaging (fMRI) to evaluate insomnia-related alterations in functional connectivity (FC). A recent review shows some divergence in the results, but also highlight the relevance of the salience network and show how alterations may be linked to hyperarousal and affective symptoms in insomnia disorder (ID) (Khazaie et al., 2017). It has recently been suggested that ID could be related to distributed deviations in various brain circuits related to salience processing, and that these distributed minor deficiencies may result in either increased arousal and alertness or dysfunctional inhibition (Van Someren, 2021).

Previous studies have focused on static FC, i.e., the average coactivation of brain areas over a time window of typically 5-10 min. However, we know that brain activity changes at a much faster pace (Deco, Cruzat, & Kringelbach, 2019; Kringelbach & Deco, 2020). Specifically, large-scale brain networks have been found to transition in time in a non-random fashion that is participant-specific and related to behavioural traits (Vidaurre, Smith, & Woolrich, 2017). So far, only one study has evaluated the temporal dimension of insomnia related alterations of brain networks. Using a sliding window approach, Wei and colleagues found reduced variability between the anterior salience network and the left executive-control network in participants with ID compared to controls (Wei et al., 2020). Since the study found no differences in static FC, it indicates that methods including the temporal dynamics may reveal more subtle differences not captured by traditional FC methods focusing only on the spatial dimension.

In this study, we used a data-driven approach to investigate the dynamic aspects of brain state transitions in ID compared to matched controls with no sleep problems. We evaluated the dynamic aspects of whole-brain activity using Hidden Markov Modelling and found evidence of reduced transitioning between brain states in the insomnia participants. Differences in the specific brain states were investigated and the findings were contextualised in relation to canonical resting-state networks.

## Methods

### Participants

We included 29 participants including 15 adults meeting the DSM-5 criteria for ID, and 14 age- and sex-matched controls with no sleep complaints. Participants with ID were screened for other sleep disorders and underwent 1 night of ambulant polysomnography. Participants were excluded if they had more than mild symptoms of other sleep disorders such as circadian rhythm disorder, sleep-related movement disorder or sleep-disordered breathing. Exclusion criteria for all participants were (a) use of psychotropic or hypnotic medications, (b) sleep-disruptive medical disorder, (c) psychiatric disorder (d) alcohol or substance abuse, and (e) MRI contraindications. All participants completed the Pittsburgh Sleep Quality Index (Buysse, Reynolds III, Monk, Berman, & Kupfer, 1989) and the Insomnia Severity Index (Bastien, Vallieres, & Morin, 2001) to evaluate the degree of sleep difficulties and insomnia severity. Participants signed informed consent in accordance with the declaration of Helsinki, and the study was approved by the ethical committee of the Central Denmark Region.

### Questionnaires

The Pittsburgh Sleep Quality Index (PSQI) was used to evaluate the degree of sleep difficulties in all participants. The PSQI is a self-report questionnaire with 19 items. The score ranges from 0 to 21 with higher scores indicating more sleep problems. A score >5 separates good sleepers from poor sleepers (Buysse, Reynolds III, Monk, Berman, & Kupfer, 1989). Insomnia severity was assessed using the Insomnia Severity Index (ISI). The ISI is a 7-item self-report questionnaire evaluating the nature and severity of insomnia. Each item is rated on a five-point scale yielding a total score between 0 and 28. Higher scores indicate more severe insomnia (Bastien, Vallieres, & Morin, 2001). Both the PSQI and ISI are commonly used in sleep research and have good psychometric properties (Bastien et al., 2001; Buysse et al., 1989; Morin, Belleville, Bélanger, & Ivers, 2011).

### Imaging protocol

For the resting-state fMRI, participants were instructed to keep their eyes open and stay awake during the 10 minutes scan. Wakefulness was validated verbally after the scan. Participants were scanned on a Siemens Trio 3-tesla MRI scanner with a 12-channel head coil at Aarhus University Hospital, Denmark. Resting-state functional images were acquired using whole-brain echo planar imaging with the following parameters: repetition time = 3,500 ms, echo time = 28 ms, voxel size = 2.0 × 2.0 × 2.4 mm^3^, slice thickness = 2.4 mm, field of view = 192 × 192 × 120 (RL x AP x FH), pixel bandwidth = 1796 Hz/Px and flip angle = 90°. Additionally, the T1-weighted images were acquired with the following parameters: repetition time = 2,000 ms, echo time = 3.7 ms, voxel size = 1 × 1 × 1 mm^3^, slice thickness = 1 mm, matrix size = 256 × 256, field of view = 256 × 256, pixel bandwidth = 150 Hz/Px, and flip angle = 9°.

### Image pre-processing

We pre-processed the fMRI data using the FSL MELODIC (www.fmrib.ox.ac.uk/fsl). This included 1) motion correction within each scanning session using FSL’s MCFLIRT (Jenkinson, Bannister, Brady, & Smith, 2002), 2) a high-pass temporal filter applied to the EPI images with a cutoff of periods longer than 100 s, 3) spatial smoothing of the EPI with a Gaussian kernel with FWHM of 4 mm, 4) slice-timing correction using Fourier-space time-series phase-shifting, 5) grand-mean intensity normalization of all images using a single multiplicative factor, and 6) decomposition of the session-specific voxel data using Probabilistic Independent Component Analysis (Beckmann & Smith, 2004) for the purpose of labelling and removing artefactual parts of the data (see below). Transformation matrices between the EPI images and MNI space were obtained in FSL via the individual T1 scans. Registrations between the EPI data and the T1 scan, and subsequently the T1 scan and the MNI152 brain were performed using FLIRT with the BBR algorithm and an affine transformation respectively. These two separate transformations were concatenated to obtain a registration between the EPI data and MNI. In cases where, according to visual inspection, the BBR provided poor registrations between EPI and T1, a simpler 6 degrees-of-freedom rigid body transformation was employed. Brain extraction was performed, and visually evaluated, on the anatomical T1 scans prior to registrations, using FSLs Brain Extraction Tool (BET, (Smith, 2002)). Artefacts were removed using ICA. We followed the approach of Griffanti and colleagues (Griffanti et al., 2017) with two researchers (K.V.J. and A.S.) independently doing a manual classification of the ICA components resulting from the MELODIC processing. Each component was labelled as either brain or non-brain signal based on its spatial distribution of activity, its time series and power spectra, according to the recommendations by Griffanti and colleagues (Griffanti et al., 2017). The independent manual classifications were subsequently compared, and the inter-rater agreement was 82%. Disagreements were discussed to reach consensus. The consensus-based labelling was used as input to a regression step. Using the FSL software FIX (Salimi-Khorshidi et al., 2014), the signal corresponding to the non-brain components, together with the motion confounds (24 regressors) derived from the motion parameters identified with MCFLIRT (see above) were regressed out of the EPI data.

### Data analysis

The analysis flow of the study is outlined in Figure 1.

**Figure 1.**
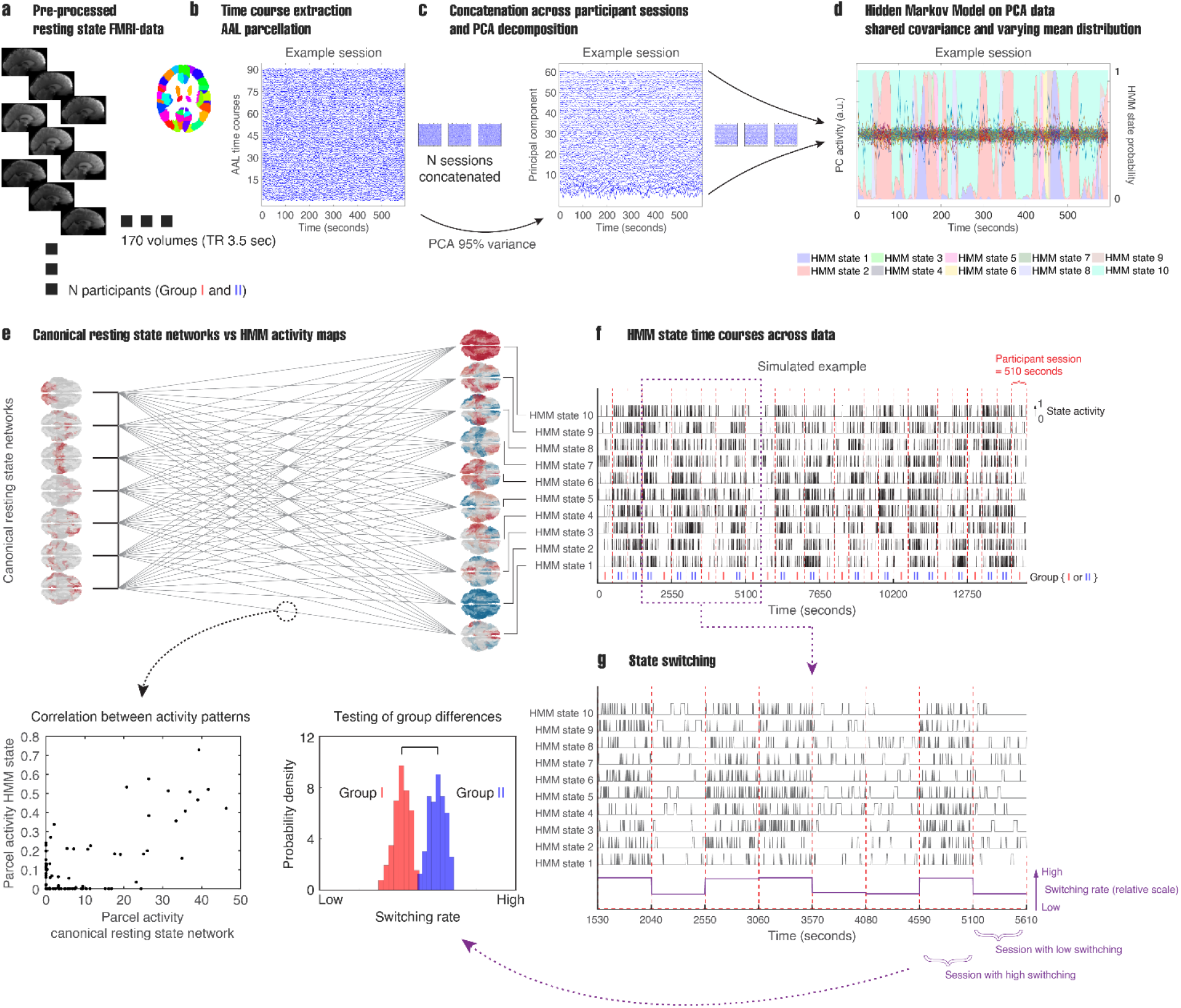
Analysis steps of the study. **a)** The preprocessed data **b)** was co-registered to the AAL template in MNI space before **c)** PCA decomposition. **d)** We applied a Hidden Markov Model (HMM) to estimate whole-brain states across all participants. **e)** The data-driven HMM states were compared to canonical resting state networks, and **f)** the estimated time courses of each HMM state were extracted. **g)** To evaluate the overall dynamics of the HMM states, we considered the switching rate, i.e. the average rate of transitioning from one HMM state to any of the other states across the recording session. Finally, we tested for group differences between the insomnia participants and matched controls.

#### AAL template

The pre-processed voxel-wise fMRI data was co-registered to the Automated Anatomical Labelling (AAL, (Tzourio-Mazoyer et al., 2002)) atlas in MNI space using the transformation matrices produced by FLIRT (see Image pre-processing). For each recording session 90 cortical and sub-cortical time-courses were derived by taking the mean across voxels within each region-of-interest (ROI). Following this step each participant was represented by data of dimensions 90-by-170 corresponding to AAL regions and the approx. 10 minutes of resting-state scanning at a TR of 3.5 seconds.

#### Static functional connectivity

As a first step, we calculated static functional connectivity measures. For each participant we used the 90-by-170 ROI time courses to construct a static functional connectivity (sFC) matrix. sFC was defined as the linear correlation between each pair of ROI time courses yielding a 90-by-90 FC matrix across the entire resting-state scan. To promote normality each participant-specific sFC matrix was Fisher z-transformed using the built-in MATLAB function *atanh*.*m*. We tested for differences in sFC between the two groups using network-based statistics (NBS, (Zalesky, Fornito, & Bullmore, 2010)) (see supplementary material). The NBS analyses showed that compared to matched controls with no sleep complaints, participants with ID were characterised by a network of increased sFC (t-threshold = 3.5, p = 0.0408). The network included 13 regions and 13 connections between them (supplementary Figure S1). This group difference in sFC provided a solid fundament for the further analyses evaluating the temporal dynamics of the brain activity.

#### Hidden Markov Modelling

To derive the spatiotemporal dynamics of the fMRI signal, we applied a Hidden Markov Model (HMM, (Vidaurre et al., 2018; Vidaurre et al., 2017)) to the AAL time courses from the ID and control population together. Prior to applying the HMM, we standardised the 90-by-170 AAL time courses within participants to have zero mean and standard deviations of 1. These standardised time-courses were then concatenated in time, leaving a data matrix of dimensions 90-by-(170 x N_participants_). This was submitted to a principal component analysis (PCA), which was used to reduce the dimensionality of the data, which is helpful to avoid issues in the HMM estimation (Ahrends et al., 2022; Charquero-Ballester et al., 2022; Stevner et al., 2019). We tested the HMM on a varying number of PCA components corresponding to different amounts of explained variance in the data. At a given PCA dimensionality an HMM was fitted with the following parameters. The observation model used was a multivariate Gaussian distribution. Unlike previous demonstrations of the HMM on fMRI data (Baldassano et al., 2017; Shappell, Caffo, Pekar, & Lindquist, 2019; Stevner et al., 2019; Vidaurre et al., 2017), we chose to restrict the model such that only the mean distributions were allowed to vary between states. In this way, the covariances were shared across states, and as such the model was sensitive mainly to changes in mean activation in the 90 AAL areas, and less sensitive to changes in functional connectivity. This was because the number of time points per subjects was relatively short to obtain a reliable estimation of time-varying FC (Ahrends et al., 2022; Alonso & Vidaurre, 2023). Fixing the covariance and allowing only the mean to vary across time reduces the complexity of the model, allowing for better model fitting. We ran the HMM for each PCA dimensionality with several numbers of states ranging between 4 and 40 in steps of 1. Ultimately, we opted for the solution with a PCA dimensionality of 95% and 10 states. We considered this a good trade-off between detail and robustness in terms of the number of states and a way of not discarding too much of the data through the PCA decomposition. Due to the lack of an objective criteria for choosing these parameters we chose to investigate the main outcome (i.e. switching) across varying PCA dimensionalities and numbers of HMM states.

#### Permutation test to compare groups (fractional occupancy)

Once an HMM was fitted to the pre-processed AAL data, concatenated across participants, a series of direct and derived outputs were produced. Each HMM state was characterised by its time course which defined the points in time for which a given state was active, and a mean distribution, corresponding to the modelled mean activation across the 90 AAL areas. The state time courses could be used to assess how the HMM states’ temporal aspects related to ID and controls. The simplest statistic is the fractional occupancies (FO): the proportion of time that each state was active per session (Vidaurre et al., 2018). Each HMM state was thus associated with a session-specific FO, ranging between 0 and 1, and these could be compared between the groups of ID and controls. This was achieved in a non-parametric GLM framework using random permutations to test for significant differences for each HMM state. We used the HMM-MAR toolbox function hmmtest.m for this step with 1000 permutations of the design matrix. We used a significance level of p < 0.05, and differences reported on FO were not corrected for the multiple comparisons created by running a test for each state.

#### Switching rate

The fact that the HMM states were defined in time made it possible to assess their dynamics. For this we considered the transition matrix. The transition matrix was defined as a K-by-K matrix, with K being the number of HMM states, and it described the probability of transitioning from a given state to any of the remaining states. The transition matrix was thus a summary of the individual state-to-state transitions modelled by the HMM. As a measure of the overall dynamics we chose to consider the average rate of these transitions across the span of a recording session, which we refer to as switching rate. To estimate this, we regarded the HMM state time-courses which were output by the model as an [n_states_ x time] matrix of values between 0 and 1, with each column representing the probability of a given state being active. To compute the transitions, we used the Viterbi path, a hard-assignment version of the state time courses where each state is either active or non-active, instead of having a probability of activation (Lou, 1995). This way, it was straightforward to count the points in time where a state change occurred and divide this by the duration of the time period in question (i.e. the length of the recording session).

We went on to compare the overall switching across the groups of ID and controls. Differences in overall switching between ID and controls were tested using the HMM-MAR toolbox function hmmtest.m. 1000 permutations of the design matrix was used to create a non-parametric null for the unpaired two-sample t-test. No multiple comparisons were involved in this test, and p-values < 0.05 were reported.

#### Correlation with clinical measures

To investigate how the switching rates relate to the clinical characteristics of the participants, we used linear regression to model the switching rate as a function of insomnia severity scores. This also allowed us to control for potential confounding effects of age, gender and movement parameters.

## Results

### Participant characteristics

The study population consisted of 29 adults. Fifteen participants fulfilled the DSM-5 criteria for insomnia disorder and 14 were matched controls with no sleep complaints. The two groups did not differ in age and gender but were clearly distinct with regard to sleep quality as reflected in the Pittsburgh Sleep Quality Index (PSQI) and the Insomnia Severity Index (ISI) (Table 1). The PSQI scores ranged from 2 to 5 in the control group and from 8 to 19 in the ID group indicating sleep difficulties only in the ID group. The ISI scores in the control group ranged from 0 to 9 in the control group and from 9 to 24 in the ID group.

**Table 1.**
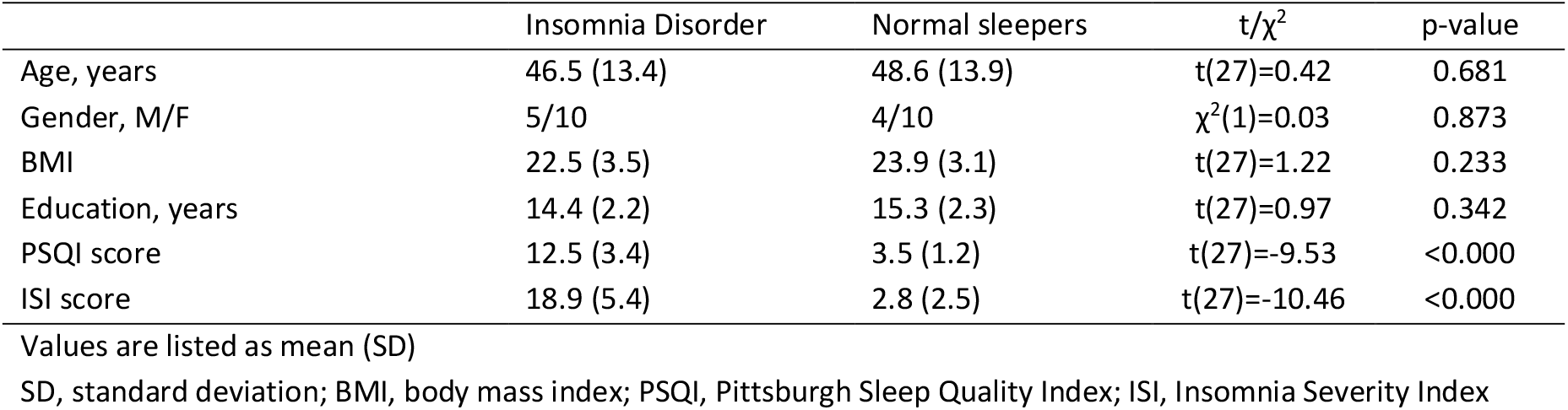
Participant characteristics.

### Between-group differences in spatiotemporal dynamics of brain states

In order to estimate the spatiotemporal organisation of whole-brain networks in ID compared to matched controls, we used the HMM on the resting-state fMRI data of all participants. The HMM estimated the probability of a given number of brain states being active at a given point in time by including the mean level of activity in each brain region. The HMM was endowed with 10 states and identified recurring whole-brain states with associated time-courses for each state (Figure 2).

**Figure 2.**
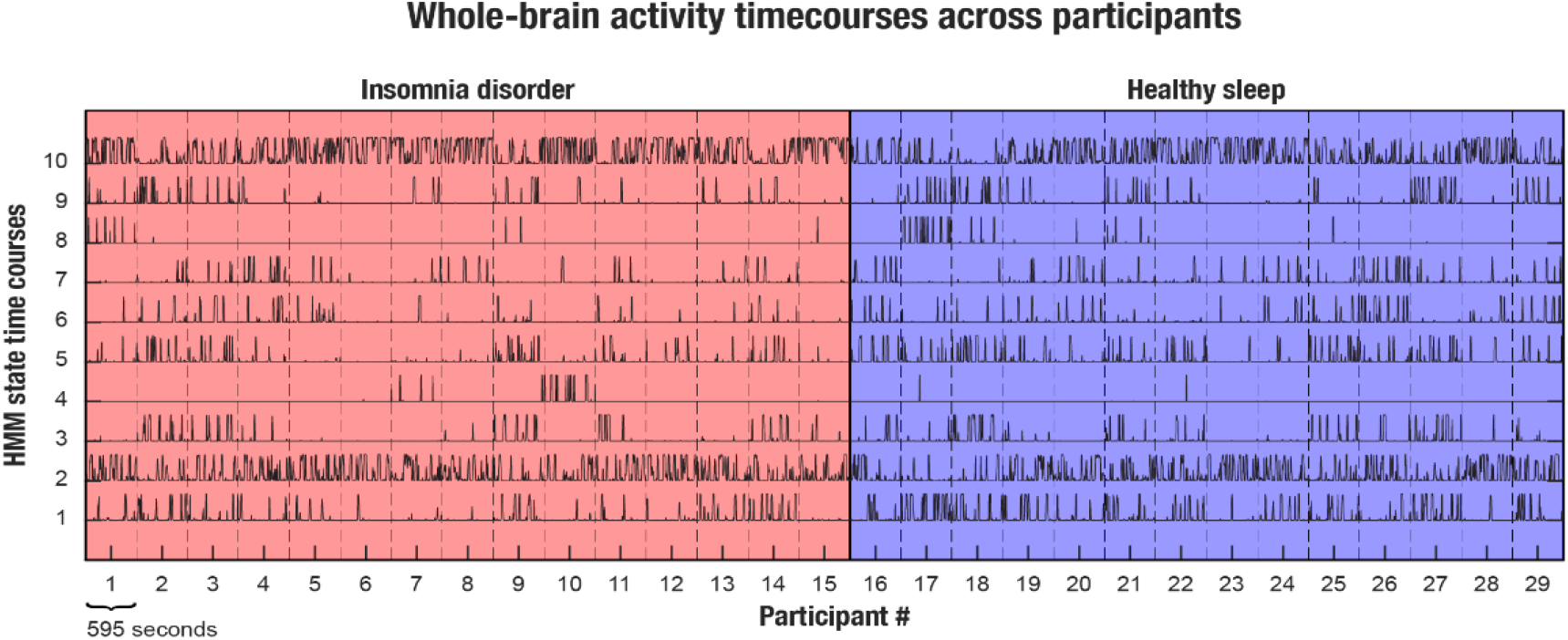
HMM time-courses for insomnia participants and good sleeper controls. Along the x-axis the participants are listed divided by the dotted lines. Insomnia participants are marked in red and good sleeper controls are marked in blue. Along the y-axis each HMM state is listed with the corresponding time-courses. The time-courses reflect that HMM states #2 and #10 are most prevalent in both groups whereas states #1 and #5 are less frequent in insomnia participants. Instead, insomnia participants tend to maintain HMM state #10 for longer time than good sleeper controls.

The switching rate of the ID participants was significantly lower than controls (p < 0.05), reflecting fewer transitions between brain states (Figure 3a). In addition, we looked at fractional occupancy, defined as the temporal proportion of a measurement, in which an HMM state is active (Vidaurre et al., 2018). The results showed a significant difference between groups in the fractional occupancy of three whole-brain states (Figure 3b). Compared to controls, ID participants spent significantly less time in two HMM brain states - one network resembling the default mode network (DMN) and the other a network of right-lateralized frontal and parietal regions. Instead, ID participants showed increased prevalence of a global activation state (Figure 3c). See supplementary Figure S2 for a visualization of all ten states.

**Figure 3.**
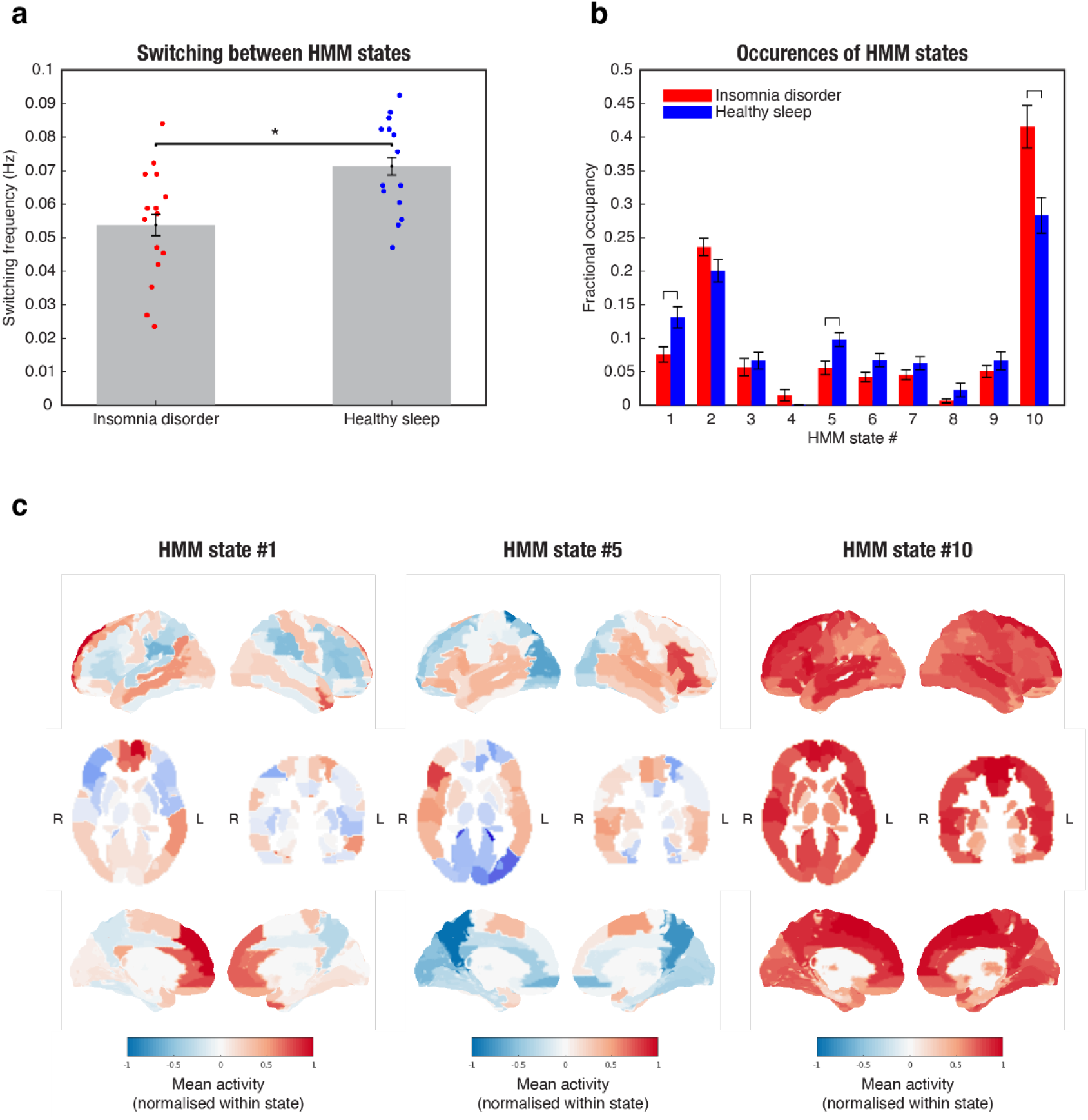
Switching, fractional occupancy and mean activity of HMM states that differ between ID and HS. **a)** the mean switching rate of the insomnia participants (red) was significantly lower than the healthy sleeper controls (blue). Significance level *p* < 0.05. **b)** the fractional occupancy for the HMM states #1 and #5 was significantly lower in the insomnia group (red) compared to controls (blue), whereas HMM state #10 was significantly more prevalent in the insomnia group compared to controls. Significance level *p* < 0.05. **c)** the mean activity of the three HMM states that were significantly different between the groups. HMM state #1 is characterized by increased activity in the dorsolateral and medial prefrontal cortex as well as in the angular gyrus and posterior cingulate. HMM state #5 show increased activity in fronto-parietal regions with pronounced activity in the ventrolateral prefrontal cortex in the right hemisphere. HMM state #10 is characterized by increased levels of activity across the entire brain (see Supplementary Figure S2 for all HMM states).

The lower switching rate in the insomnia group was found to be consistent across PCA dimensionalities and number of states of the HMM, indicating that the findings are robust and not dependent on specific HMM parameters (Supplementary Figure S3).

To evaluate the correspondence between the HMM states and canonical resting-state networks (Yeo et al., 2011), we did two correlation matrices (Figure 4). These results confirm a very high degree overlap between HMM state 1 and the DMN. HMM state 5 is positively correlated with the frontoparietal network, but also has a high correlation with the ventral attention network.

**Figure 4.**
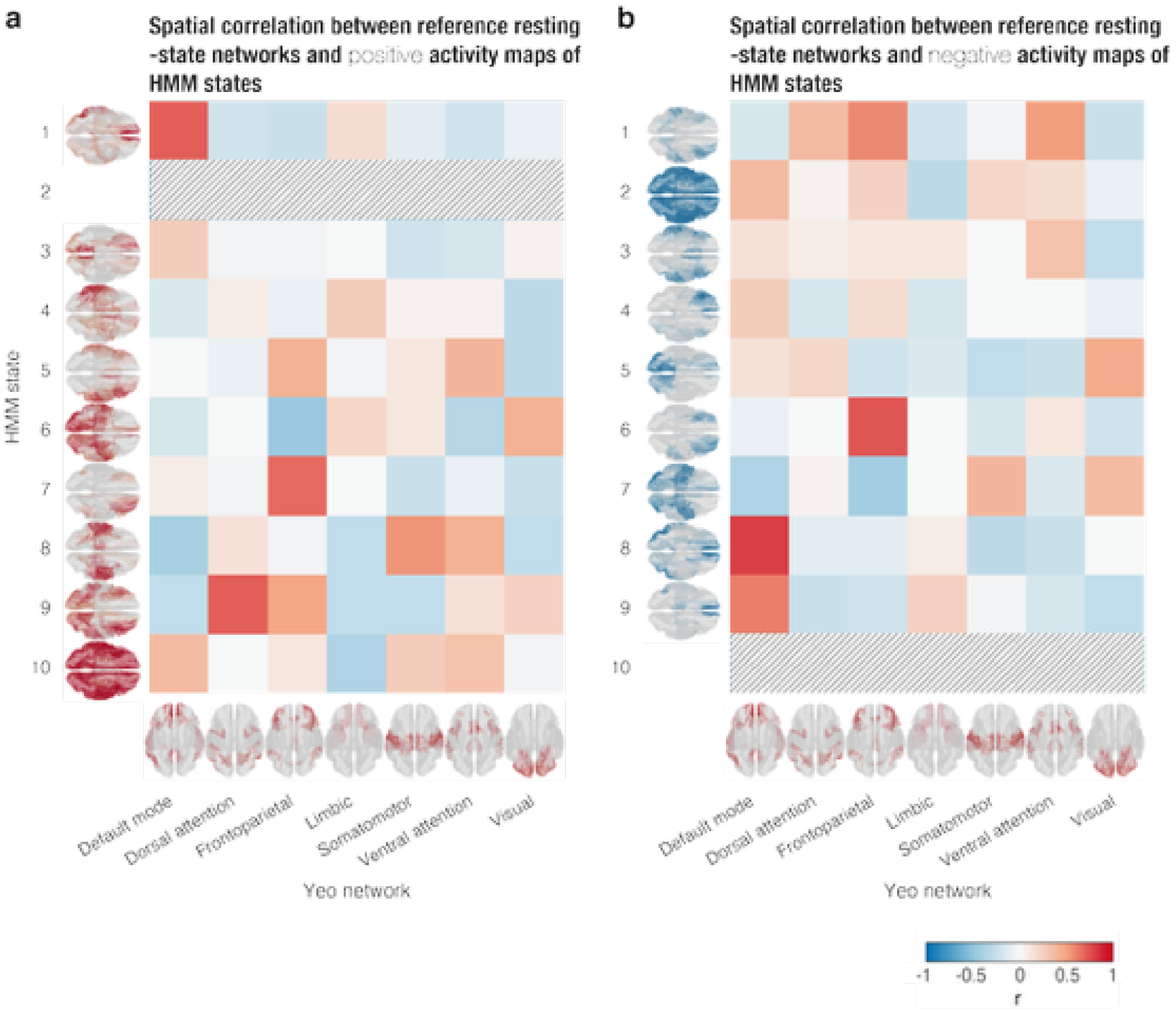
Spatial correlation between the HMM state maps and canonical resting states from Yeo et al. (2011). Section **a)** reflects the positive activity maps. HMM state #2 includes only deactivation and is therefore left blank. Similarly, section **b)** reflects the negative activity maps, i.e. deactivation, and here, HMM state #10 is left blank because it includes no deactivation. Note the strong positive correlation with the default mode network in **a)** and HMM state #1.

### Correlation with insomnia severity

We used linear regression to investigate the relationship between switching rates and insomnia severity measured with the insomnia severity index while controlling for age and sex. The results showed a significant negative linear relationship between switching rate and insomnia severity in all participants (F(4, 25) = 5.77, *p* = 0.0038, R2 = 0.409). Only insomnia severity was a significant predictor of switching rate (*p* = 0.0016). Similar results were found when including motion and group*insomnia severity interaction in the model (see Figure 4 and supplementary Table S1). This means that higher insomnia severity scores were associated with lower switching rate (Figure 5). There were no significant correlations when looking at the groups separately.

**Figure 5.**
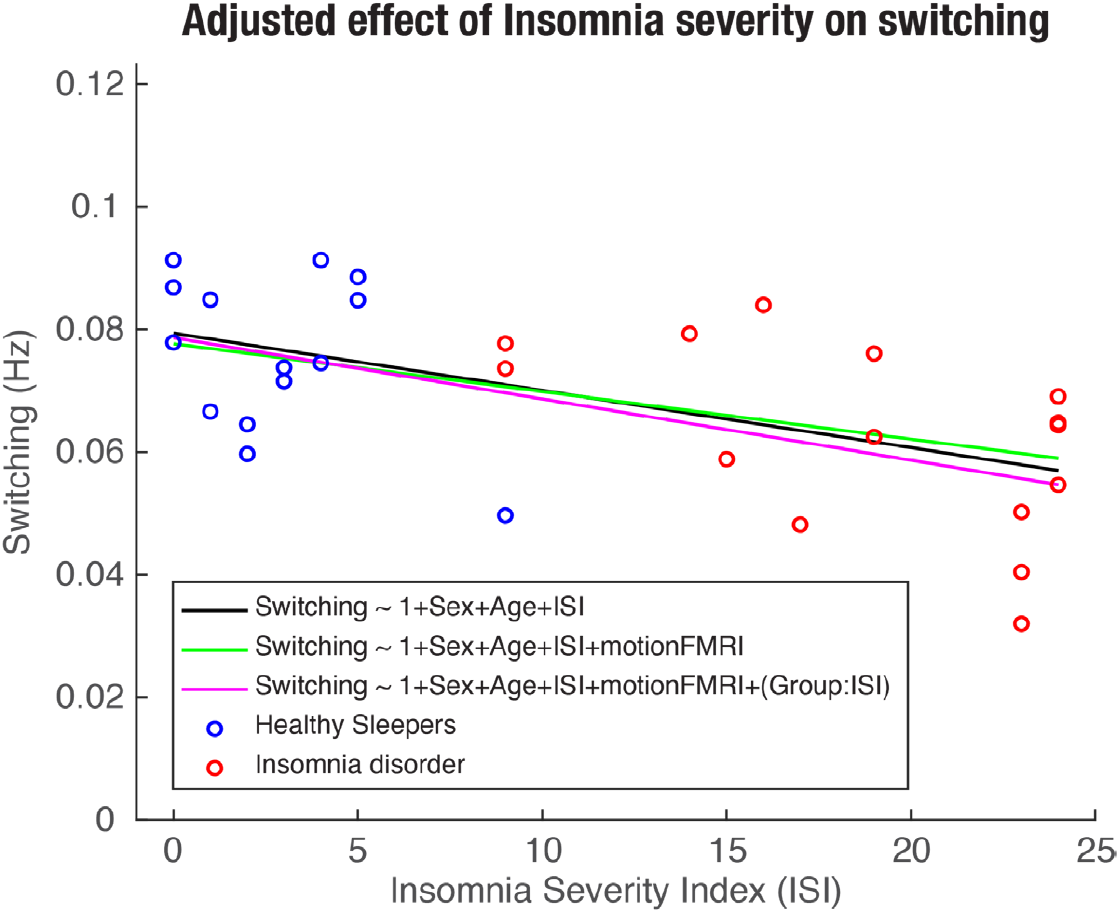
The linear regression model showed a significant relationship between insomnia severity and the rate with which between brain states transition from one to another. Higher insomnia scores are related to lower switching rate across all participants.

## Discussion

We found reduced switching between brain states in participants with insomnia disorder (ID) compared to control participants with no sleep problems. Using Hidden Markov Modelling (HMM) as a data-driven approach on resting-state fMRI data we saw that on average, ID participants transitioned between brain states less often than controls. Specifically, the fractional occupancy of the HMM revealed that compared to controls, insomnia participants spent more time in a global activation state and less time in two specific states including a network of right frontal and parietal regions and the default mode network. These results expand our knowledge on the neural underpinnings of ID and suggest reduced flexibility in the temporal dynamics of brain activity in insomnia disorder.

One previous study has examined the temporal dynamics of brain networks in ID. This study focused on established resting-state networks using a sliding-windows approach and found less functional connectivity variability between the anterior salience network and the left executive-control network, as well as a trend towards reduced variability between the anterior salience network and the dorsal default-mode network (Wei et al., 2020). This is in line with our main result showing reduced switching between brain states in insomnia compared to controls. The approach of the two studies differ in the methods used to analyse the time-varying aspects of brain activity (see (Ahrends & Vidaurre, 2023) for an overview). Furthermore, the study by Wei and colleagues compared the functional connectivity of canonical resting-state networks, whereas our study took a data driven approach comparing the whole-brain states revealed by the HMM based on mean brain activity. Nevertheless, the results reflect the same dynamic in the neural activity over time, i.e. fewer transitions between whole brain states in ID during rest. Hereby, the results highlight the importance of using analyses that include the time-varying aspect of brain activity in order to advance our understanding of the neural fundament of ID and thereby inform novel treatment approaches.

When looking at the specific brain states, we found that ID participants spent more time in a global activation state and less time in specific states such as the network of right frontal and parietal regions and the default-mode network. The higher prevalence of this global activation state could be interpreted as an expression of cortical hyperarousal in ID. Hyperarousal is often seen as a core feature of insomnia reflected in both physiological, cortical and cognitive measures (Riemann et al., 2015; Van Someren, 2021).

Hyperarousal in ID is considered a 24-hour condition which matches well with the fact that our participants were scanned at daytime. However, we must note that this ‘global signal’ is commonly found in resting state fMRI studies of functional connectivity, and its role and neurophysiological origin is discussed (Bennett, Farnell, Gibson, & Lagopoulos, 2016; Cabral et al., 2017; Vohryzek, Deco, Cessac, Kringelbach, & Cabral, 2020). Therefore, the interpretation of this state must be carefully considered. In this study, we included mean activation, but not covariance in the HMM analysis, and as such, the results are not directly comparable to functional connectivity studies.

In contrast to the higher prevalence of the global activation brain state, participants with insomnia had significantly less activation of the HMM brain state corresponding to the default mode network (see Figure 4). The default mode network is primarily related to internally oriented mental processes such as mind wandering, remembering and future planning (Buckner, Andrews-Hanna, & Schacter, 2008; Raichle, 2015; Smallwood et al., 2021). DMN activation has previously been linked to hyperarousal reduction in PTSD patients (King, Angstadt, Sripada, & Liberzon, 2017), and the reduced DMN activity seen in our insomnia participants could be interpreted as a result of hyperarousal. Altered DMN activity has previously been related to ID (Kay & Buysse, 2017; Khazaie et al., 2017; Marques, Gomes, Caetano, & Castelo-Branco, 2018). For example, Nie and colleagues evaluated functional connectivity (FC) between subregions of the DMN in participants with primary insomnia compared to good sleeper controls, and found significantly decreased FC between DMN subregions in insomnia participants (Nie et al., 2015). Interestingly, previous research investigating whole brain transitioning in sleep has suggested that increased DMN activity may serve as a gate for sleep initiation (Stevner et al., 2019). This study used combined EEG and fMRI to characterise the brain dynamics underlying sleep and sleep state transitioning. The results showed that the key trajectory from wakefulness to sleep went through the DMN. Related to our results, this suggests that reduced DMN activation at rest in insomnia participants may underlie difficulties transitioning from wakefulness to sleep.

### Limitations

The findings of this study are limited by the relatively small sample size, and they should be replicated in a larger dataset. However, the two groups are clearly distinct in terms of clinical measures while at the same time well-matched. The insomnia participants are clearly defined, and other underlying sleep disorders are ruled out by the initial polysomnography assessment. Another limitation is the cross-sectional design that does not enable us to make causal conclusions. Based on this study, we cannot conclude if the altered brain activity is a predisposition for ID or if it is the result of the disorder. Future studies would benefit from a longitudinal design to disentangle the causal relationship.

## Conclusion

This is one of the first studies investigating the dynamic aspects of resting-state brain activity in insomnia disorder compared to matched controls, and the findings show fewer transitions between brain states in ID. These results suggest reduced flexibility in the neural dynamics of whole-brain activity in ID and highlight the importance of using methods that account for temporal dynamics to advance our understanding of the neural foundations of a complex disorder like ID.

## Acknowledgements

Center for Music in the Brain is funded by the Danish National Research Foundation (DNRF117).

## References

Ahrends, C., Stevner, A., Pervaiz, U., Kringelbach, M. L., Vuust, P., Woolrich, M. W., & Vidaurre, D. (2022). Data and model considerations for estimating time-varying functional connectivity in fMRI. NeuroImage, 252, 119026. doi:10.1016/j.neuroimage.2022.119026

Ahrends, C., & Vidaurre, D. (2023). Dynamic Functional Connectivity. arXiv preprint 2301.03408.

Alonso, S., & Vidaurre, D. (2023). Toward stability of dynamic FC estimates in neuroimaging and electrophysiology: Solutions and limits. Network Neuroscience, 7(4), 1389–1403. doi:10.1162/netn_a_00331

Aquino, G., Benz, F., Dressle, R. J., Gemignani, A., Alfì, G., Palagini, L., … Feige, B. (2024). Towards the neurobiology of insomnia: A systematic review of neuroimaging studies. Sleep Medicine Reviews, 73, 101878. doi:10.1016/j.smrv.2023.101878

Baldassano, C., Chen, J., Zadbood, A., Pillow, J. W., Hasson, U., & Norman, K. A. (2017). Discovering event structure in continuous narrative perception and memory. Neuron, 95(3), 709-721. e705.

Bastien, C. H., Vallieres, A., & Morin, C. M. (2001). Validation of the Insomnia Severity Index as an outcome measure for insomnia research. Sleep Medicine, 2(4), 297–307.

Beckmann, C. F., & Smith, S. M. (2004). Probabilistic independent component analysis for functional magnetic resonance imaging. IEEE transactions on medical imaging, 23(2), 137–152.

Bennett, M. R., Farnell, L., Gibson, W., & Lagopoulos, J. (2016). On the origins of the ‘global signal’ determined by functional magnetic resonance imaging in the resting state. Journal of Neural Engineering, 13(1), 016012. doi:10.1088/1741-2560/13/1/016012

Buckner, R. L., Andrews-Hanna, J. R., & Schacter, D. L. (2008). The brain’s default network: anatomy, function, and relevance to disease. Annals of the New York Academy of Sciences, 1124(1), 1–38.

Buysse, D. J., Reynolds III, C. F., Monk, T. H., Berman, S. R., & Kupfer, D. J. (1989). The Pittsburgh sleep quality index: A new instrument for psychiatric practice and research. Psychiatry Research, 28(2), 193–213. doi:10.1016/0165-1781(89)90047-4

Cabral, J., Vidaurre, D., Marques, P., Magalhães, R., Silva Moreira, P., Miguel Soares, J., … Kringelbach, M. L. (2017). Cognitive performance in healthy older adults relates to spontaneous switching between states of functional connectivity during rest. Scientific reports, 7(1), 5135. doi:10.1038/s41598-017-05425-7

Charquero-Ballester, M., Kleim, B., Vidaurre, D., Ruff, C., Stark, E., Tuulari, J. J., … Moseley, A. (2022). Effective psychological therapy for PTSD changes the dynamics of specific large-scale brain networks. Human brain mapping, 43(10), 3207–3220.

Deco, G., Cruzat, J., & Kringelbach, M. L. (2019). Brain songs framework used for discovering the relevant timescale of the human brain. Nature communications, 10(1), 583. doi:10.1038/s41467-018-08186-7

Griffanti, L., Douaud, G., Bijsterbosch, J., Evangelisti, S., Alfaro-Almagro, F., Glasser, M. F., … Smith, S. M. (2017). Hand classification of fMRI ICA noise components. Neuroimage, 154, 188–205. doi:10.1016/j.neuroimage.2016.12.036

Jenkinson, M., Bannister, P., Brady, M., & Smith, S. (2002). Improved Optimization for the Robust and Accurate Linear Registration and Motion Correction of Brain Images. Neuroimage, 17(2), 825–841. doi:10.1006/nimg.2002.1132

Jespersen, K. V., Stevner, A., Fernandes, H., Sørensen, S. D., Van Someren, E., Kringelbach, M., & Vuust, P. (2020). Reduced structural connectivity in Insomnia Disorder. Journal of Sleep Research, 29(1), e12901.

Kay, D. B., & Buysse, D. J. (2017). Hyperarousal and Beyond: New Insights to the Pathophysiology of Insomnia Disorder through Functional Neuroimaging Studies. Brain Sciences, 7(3), 23. doi:10.3390/brainsci7030023

Khazaie, H., Veronese, M., Noori, K., Emamian, F., Zarei, M., Ashkan, K., … Morrell, M. J. (2017). Functional reorganization in obstructive sleep apnoea and insomnia: a systematic review of the resting-state fMRI. Neuroscience & Biobehavioral Reviews, 77, 219–231.

King, A., Angstadt, M., Sripada, C., & Liberzon, I. (2017). Increased Default Mode Network (DMN) Connectivity with Attention Networks with a Mindfulness-Based Intervention for PTSD: Seed and Whole Brain Connectomics Analyses. Biological Psychiatry, 81(10), S43–S44.

Kringelbach, M. L., & Deco, G. (2020). Brain States and Transitions: Insights from Computational Neuroscience. Cell Reports, 32(10), 108128. doi:10.1016/j.celrep.2020.108128

Lou, H.-L. (1995). Implementing the Viterbi algorithm. IEEE Signal Processing Magazine, 12(5), 42–52.

Marques, D. R., Gomes, A. A., Caetano, G., & Castelo-Branco, M. (2018). Insomnia Disorder and Brain’s Default-Mode Network. Current Neurology and Neuroscience reports, 18(8), 45. doi:10.1007/s11910-018-0861-3

Morin, C. M., Belleville, G., Bélanger, L., & Ivers, H. (2011). The Insomnia Severity Index: psychometric indicators to detect insomnia cases and evaluate treatment response. Sleep, 34(5), 601–608.

Nie, X., Shao, Y., Liu, S.-y., Li, H.-j., Wan, A.-l., Nie, S., … Dai, X.-j. (2015). Functional connectivity of paired default mode network subregions in primary insomnia. Neuropsychiatric Disease and Treatment, 11, 3085.

Ohayon, M. M. (2002). Epidemiology of insomnia: what we know and what we still need to learn. Sleep Medicine Reviews, 6(2), 97–111. doi:10.1053/smrv.2002.0186

Raichle, M. E. (2015). The brain’s default mode network. Annual review of neuroscience, 38, 433–447.

Riemann, D., Baglioni, C., Bassetti, C., Bjorvatn, B., Dolenc Groselj, L., Ellis, J. G., … Spiegelhalder, K. (2017). European guideline for the diagnosis and treatment of insomnia. Journal of Sleep Research. doi:10.1111/jsr.12594

Riemann, D., Nissen, C., Palagini, L., Otte, A., Perlis, M. L., & Spiegelhalder, K. (2015). The neurobiology, investigation, and treatment of chronic insomnia. The Lancet Neurology, 14(5), 547–558.

Salimi-Khorshidi, G., Douaud, G., Beckmann, C. F., Glasser, M. F., Griffanti, L., & Smith, S. M. (2014). Automatic denoising of functional MRI data: combining independent component analysis and hierarchical fusion of classifiers. NeuroImage, 90, 449–468.

Shappell, H., Caffo, B. S., Pekar, J. J., & Lindquist, M. A. (2019). Improved state change estimation in dynamic functional connectivity using hidden semi-Markov models. NeuroImage, 191, 243–257.

Smallwood, J., Bernhardt, B. C., Leech, R., Bzdok, D., Jefferies, E., & Margulies, D. S. (2021). The default mode network in cognition: a topographical perspective. Nature Reviews Neuroscience, 22(8), 503–513. doi:10.1038/s41583-021-00474-4

Smith, S. M. (2002). Fast robust automated brain extraction. Hum Brain Mapp, 17(3), 143–155.

Stevner, A., Vidaurre, D., Cabral, J., Rapuano, K., Nielsen, S. F. V., Tagliazucchi, E., … Woolrich, M. (2019). Discovery of key whole-brain transitions and dynamics during human wakefulness and non-REM sleep. Nature communications, 10(1), 1035.

Tahmasian, M., Noori, K., Samea, F., Zarei, M., Spiegelhalder, K., Eickhoff, S. B., … Eickhoff, C. R. (2018). A Lack of consistent brain alterations in insomnia disorder: an activation likelihood estimation meta-analysis. Sleep Medicine Reviews.

Tzourio-Mazoyer, N., Landeau, B., Papathanassiou, D., Crivello, F., Etard, O., Delcroix, N., … Joliot, M. (2002). Automated anatomical labeling of activations in SPM using a macroscopic anatomical parcellation of the MNI MRI single-subject brain. NeuroImage, 15(1), 273–289.

Van Someren, E. J. (2021). Brain mechanisms of insomnia: new perspectives on causes and consequences. Physiological Reviews, 101(3), 995–1046.

Vidaurre, D., Abeysuriya, R., Becker, R., Quinn, A. J., Alfaro-Almagro, F., Smith, S. M., & Woolrich, M. W. (2018). Discovering dynamic brain networks from big data in rest and task. NeuroImage, 180, 646–656. doi:10.1016/j.neuroimage.2017.06.077

Vidaurre, D., Smith, S. M., & Woolrich, M. W. (2017). Brain network dynamics are hierarchically organized in time. Proceedings of the National Academy of Sciences, 114(48), 12827–12832. doi:10.1073/pnas.1705120114

Vohryzek, J., Deco, G., Cessac, B., Kringelbach, M. L., & Cabral, J. (2020). Ghost Attractors in Spontaneous Brain Activity: Recurrent Excursions Into Functionally-Relevant BOLD Phase-Locking States. Frontiers in systems neuroscience, 14. doi:10.3389/fnsys.2020.00020

Wei, Y., Leerssen, J., Wassing, R., Stoffers, D., Perrier, J., & Van Someren, E. J. (2020). Reduced dynamic functional connectivity between salience and executive brain networks in insomnia disorder. Journal of Sleep Research, 29(2), e12953.

Wittchen, H.-U., Jacobi, F., Rehm, J., Gustavsson, A., Svensson, M., Jönsson, B., … Faravelli, C. (2011). The size and burden of mental disorders and other disorders of the brain in Europe 2010. European Neuropsychopharmacology, 21(9), 655–679.

Yeo, B. T., Krienen, F. M., Sepulcre, J., Sabuncu, M. R., Lashkari, D., Hollinshead, M., … Polimeni, J. R. (2011). The organization of the human cerebral cortex estimated by intrinsic functional connectivity. Journal of neurophysiology.

Zalesky, A., Fornito, A., & Bullmore, E. T. (2010). Network-based statistic: Identifying differences in brain networks. NeuroImage, 53(4), 1197–1207. doi:10.1016/j.neuroimage.2010.06.041

