## Supplementary material for "Modelling of brain dynamics reveals reduced switching between brain states in insomnia disorder - a resting state fMRI study"

*shared first-authorship

[1]Center for Music in the Brain, Department of Clinical Medicine, Aarhus University & the Royal Academy of Music Aarhus/Aalborg, Denmark

[2]Center for Functionally Integrative Neuroscience, Department of Clinical Medicine, Aarhus University

[3]Centre for Eudaimonia and Human Flourishing, Linacre College, University of Oxford, UK

[4]Department of Psychiatry, University of Oxford, UK

[5]The Netherlands Institute for Neuroscience, Amsterdam, The Netherlands

[6]Department of Integrative Neurophysiology, Center for Neurogenomics and Cognitive Research, Amsterdam Neuroscience, Vrije Universiteit Amsterdam, Amsterdam, The Netherlands

[7]Department of Psychiatry, Amsterdam Public Health Research Institute and Amsterdam Neuroscience Research Institute, Amsterdam UMC, Vrije Universiteit Amsterdam, Amsterdam, The Netherlands

**Content**

Static Functional Connectivity methods

Supplementary Figure S1 – The static functional connectivity network

Supplementary Figure S2 – The ten HMM states

Supplementary Figure S3 – Consistency in switching rate group difference between ID and HS across HMM solutions of varying PCA dimensionality and number of states.

Supplementary Table S1 – Results of the linear regression models

#### Static functional connectivity methods

**NBS to compare groups diff in static FC**. The two groups, ID and control, were tested for differences in sFC. This was done as a network-wide test, which given the multitude of unique connections in each sFC matrix, $(n_{ROI}^2 - n_{ROI})/2$, has to account for multiple comparisons. This was achieved in the framework of network-based statistics (NBS, Xx refs Zalesky 2010). In short, NBS was implemented as a General Linear Model (GLM) with a design describing a one-sided two-sample t-test between the ID and control groups. A range of t-statistic thresholds were investigated, from t>2 to t>5 in steps of 0.1, to obtain thresholded t-stat matrices on the differences between ID and controls. Clusters of connections were identified in these differential t-stat networks and characterised in terms of the sum of test statistics, i.e. the *intensity* in NBS terminology. The supra-threshold intensities of each t-stat threshold were compared to null-distributions of intensity values generated through 5000 random permutations of the design matrix. A supra-threshold network was considered significant if the probability of finding a network of similar intensity by chance was less than 5%, i.e. $p<0.05$. Importantly this method corrects for multiple comparisons on a network level, and hence can only speak to the significance of a network of FC weights in a differential t-stat matrix and not individual connections.

#### Between group difference in static functional connectivity

Compared to matched controls with no sleep complaints, the NBS analyses showed that participants with ID were characterized by a network of increased static FC (t-threshold = 3.5, p = 0.0408). The network included 13 regions and 13 connections between them (Figure S1). The region with the highest degree (i.e. number of connections within the significant difference network) is the right inferior parietal region (degree = 6), followed by the frontal inferior operculum, left caudate and the right calcarine fissure and surrounding cortex (all degree = 3) (Figure S1). The findings were robust to other NBS settings and the analyses revealed no networks of reduced static FC.

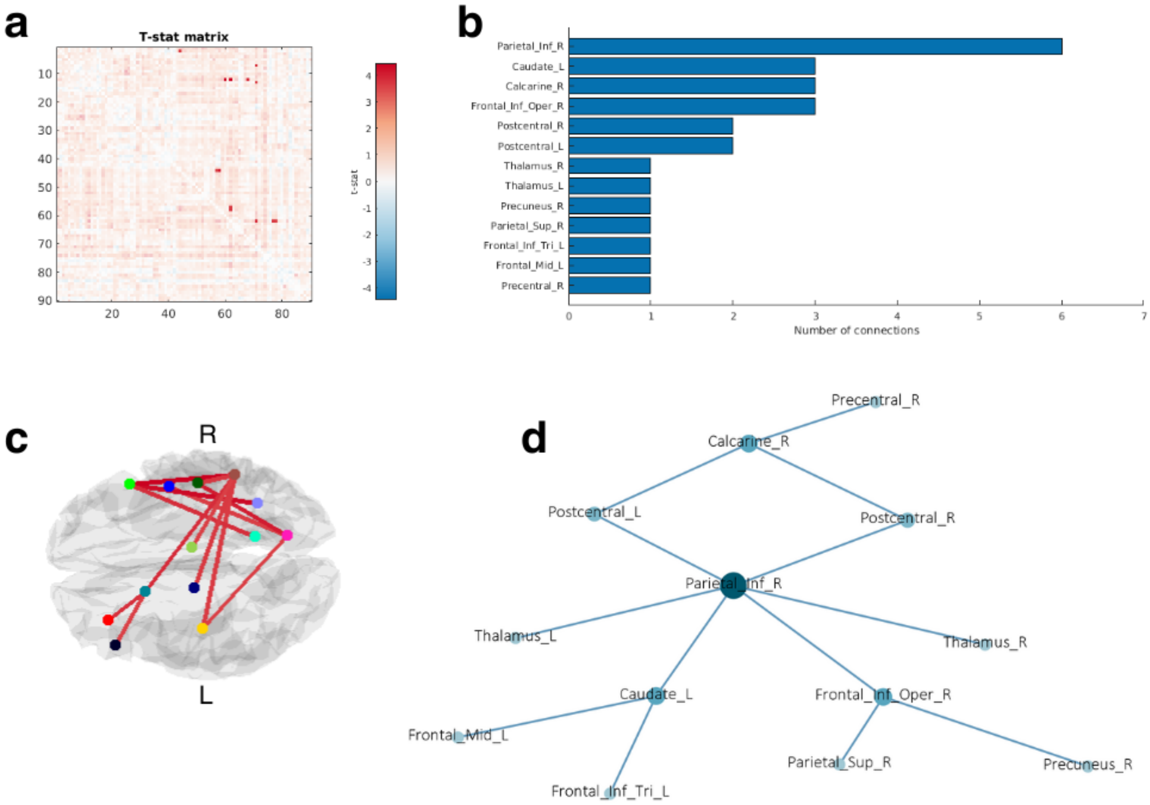


**Supplementary Figure S1** The static functional connectivity network. **a)** Correlation matrix of the T-stats. **b)** the regions included in the significant network and the number of connections for each region. **c)** Visualization of the network of increased static functional connectivity in insomnia participants compared to good sleeper controls. **d)** A two-dimensional visualization of the network highlighting the connections between each region in the network.

### The ten HMM states

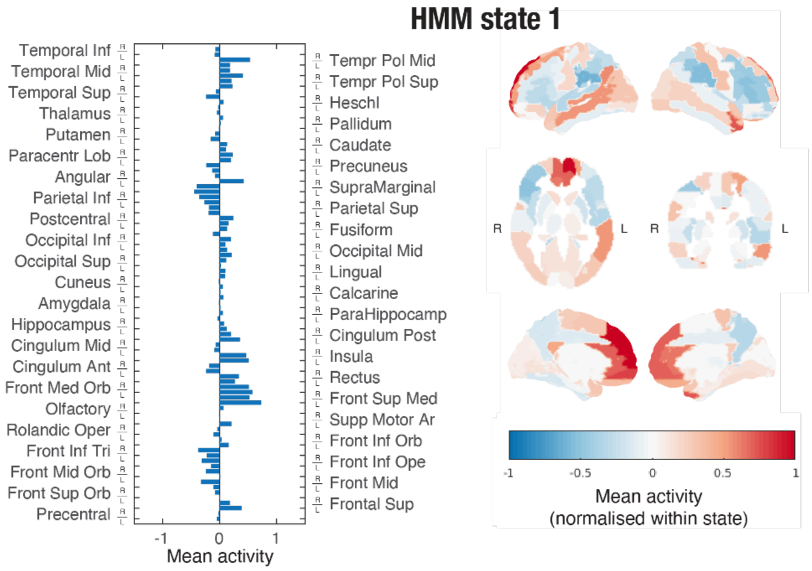


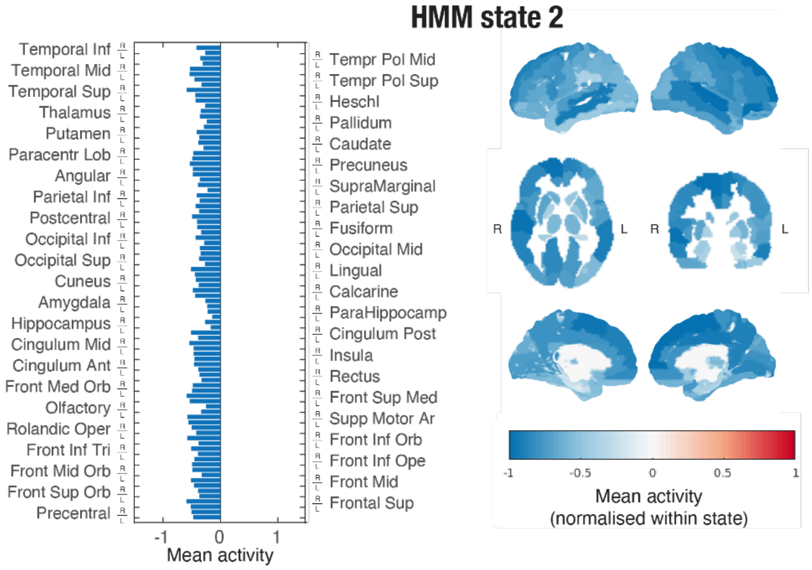


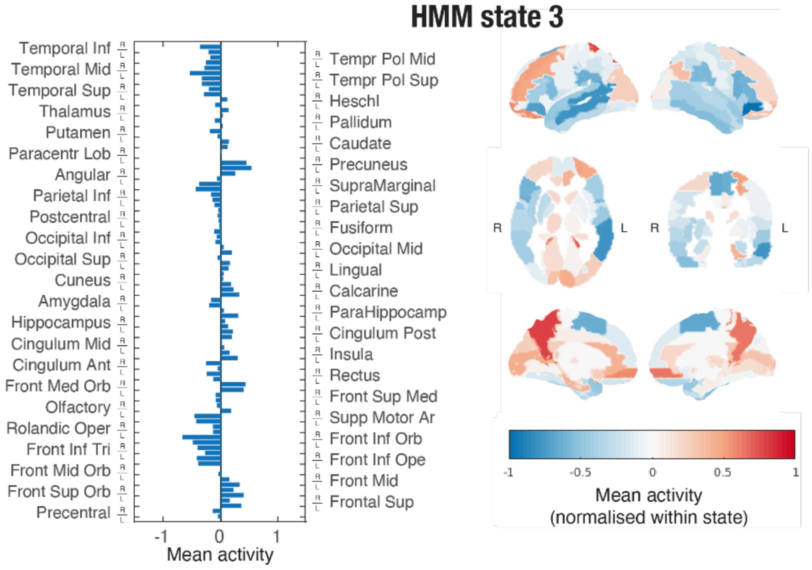

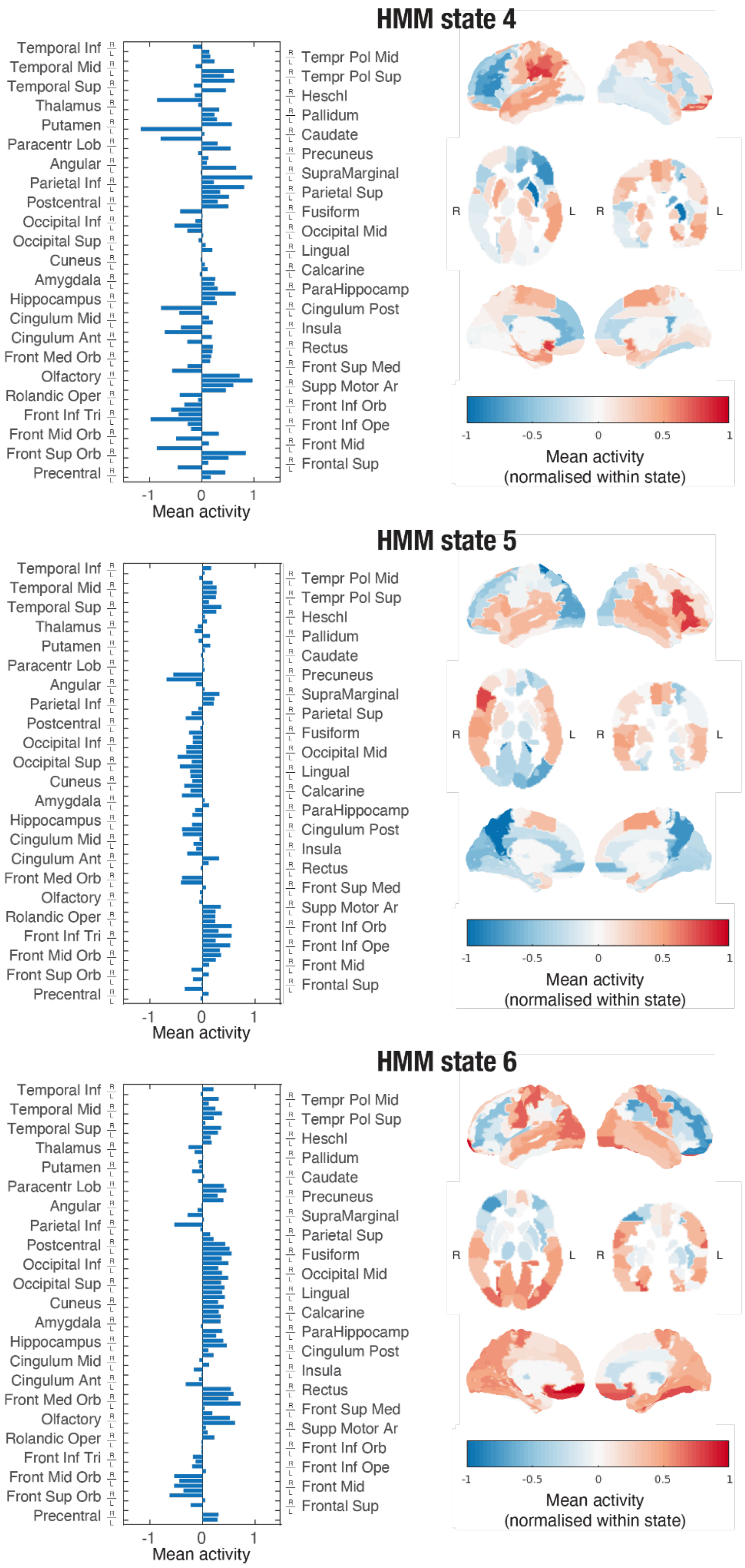
HMM state 4
Temporal Inf
Temporal Mid
Temporal Sup
Thalamus
Putamen
Paracentr Lob
Angular
Parietal Inf
Postcentral
Occipital Inf
Occipital Sup
Cuneus
Amygdala
Hippocampus
Cingulum Mid
Cingulum Ant
Front Med Orb
Olfactory
Rolandic Oper
Front Inf Tri
Front Mid Orb
Front Sup Orb
Precentral
Tempr Pol Mid
Tempr Pol Sup
Heschl
Pallidum
Caudate
Precuneus
SupraMarginal
Parietal Sup
Fusiform
Occipital Mid
Lingual
Calcarine
ParaHippocamp
Cingulum Post
Insula
Rectus
Front Sup Med
Supp Motor Ar
Front Inf Orb
Front Inf Ope
Front Mid
Frontal Sup
-1
0
1
Mean activity
R
L
R
L
-1
-0.5
0
0.5
1
Mean activity
(normalised within state)
HMM state 5
Temporal Inf
Temporal Mid
Temporal Sup
Thalamus
Putamen
Paracentr Lob
Angular
Parietal Inf
Postcentral
Occipital Inf
Occipital Sup
Cuneus
Amygdala
Hippocampus
Cingulum Mid
Cingulum Ant
Front Med Orb
Olfactory
Rolandic Oper
Front Inf Tri
Front Mid Orb
Front Sup Orb
Precentral
Tempr Pol Mid
Tempr Pol Sup
Heschl
Pallidum
Caudate
Precuneus
SupraMarginal
Parietal Sup
Fusiform
Occipital Mid
Lingual
Calcarine
ParaHippocamp
Cingulum Post
Insula
Rectus
Front Sup Med
Supp Motor Ar
Front Inf Orb
Front Inf Ope
Front Mid
Frontal Sup
-1
0
1
Mean activity
R
L
R
L
-1
-0.5
0
0.5
1
Mean activity
(normalised within state)
HMM state 6
Temporal Inf
Temporal Mid
Temporal Sup
Thalamus
Putamen
Paracentr Lob
Angular
Parietal Inf
Postcentral
Occipital Inf
Occipital Sup
Cuneus
Amygdala
Hippocampus
Cingulum Mid
Cingulum Ant
Front Med Orb
Olfactory
Rolandic Oper
Front Inf Tri
Front Mid Orb
Front Sup Orb
Precentral
Tempr Pol Mid
Tempr Pol Sup
Heschl
Pallidum
Caudate
Precuneus
SupraMarginal
Parietal Sup
Fusiform
Occipital Mid
Lingual
Calcarine
ParaHippocamp
Cingulum Post
Insula
Rectus
Front Sup Med
Supp Motor Ar
Front Inf Orb
Front Inf Ope
Front Mid
Frontal Sup
-1
0
1
Mean activity
R
L
R
L
-1
-0.5
0
0.5
1
Mean activity
(normalised within state)

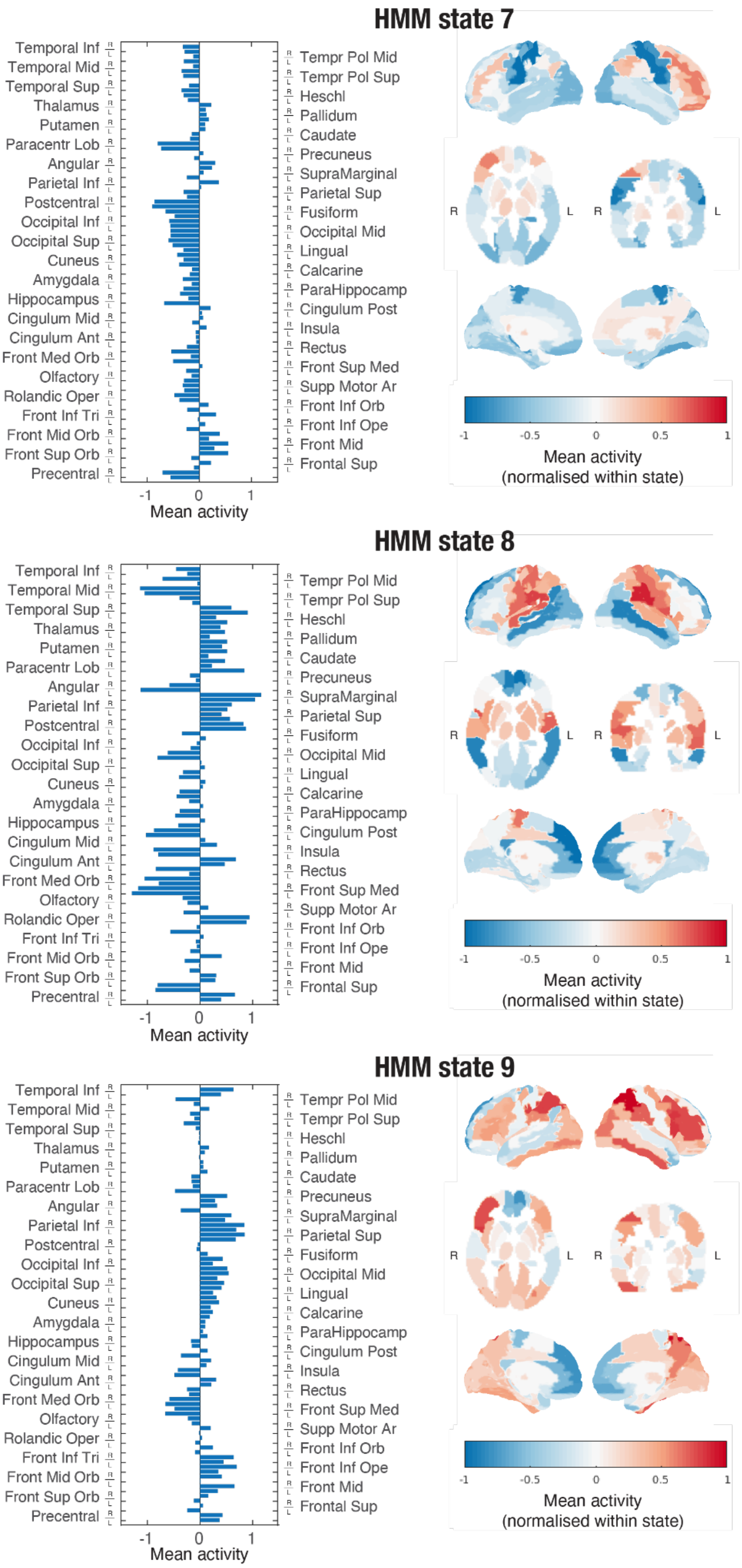
HMM state 7
Temporal Inf
Temporal Mid
Temporal Sup
Thalamus
Putamen
Paracentr Lob
Angular
Parietal Inf
Postcentral
Occipital Inf
Occipital Sup
Cuneus
Amygdala
Hippocampus
Cingulum Mid
Cingulum Ant
Front Med Orb
Olfactory
Rolandic Oper
Front Inf Tri
Front Mid Orb
Front Sup Orb
Precentral
Tempr Pol Mid
Tempr Pol Sup
Heschl
Pallidum
Caudate
Precuneus
SupraMarginal
Parietal Sup
Fusiform
Occipital Mid
Lingual
Calcarine
ParaHippocamp
Cingulum Post
Insula
Rectus
Front Sup Med
Supp Motor Ar
Front Inf Orb
Front Inf Ope
Front Mid
Frontal Sup
-1
0
1
Mean activity
R
L
R
L
-1
-0.5
0
0.5
1
Mean activity
(normalised within state)
HMM state 8
Temporal Inf
Temporal Mid
Temporal Sup
Thalamus
Putamen
Paracentr Lob
Angular
Parietal Inf
Postcentral
Occipital Inf
Occipital Sup
Cuneus
Amygdala
Hippocampus
Cingulum Mid
Cingulum Ant
Front Med Orb
Olfactory
Rolandic Oper
Front Inf Tri
Front Mid Orb
Front Sup Orb
Precentral
Tempr Pol Mid
Tempr Pol Sup
Heschl
Pallidum
Caudate
Precuneus
SupraMarginal
Parietal Sup
Fusiform
Occipital Mid
Lingual
Calcarine
ParaHippocamp
Cingulum Post
Insula
Rectus
Front Sup Med
Supp Motor Ar
Front Inf Orb
Front Inf Ope
Front Mid
Frontal Sup
-1
0
1
Mean activity
R
L
R
L
-1
-0.5
0
0.5
1
Mean activity
(normalised within state)
HMM state 9
Temporal Inf
Temporal Mid
Temporal Sup
Thalamus
Putamen
Paracentr Lob
Angular
Parietal Inf
Postcentral
Occipital Inf
Occipital Sup
Cuneus
Amygdala
Hippocampus
Cingulum Mid
Cingulum Ant
Front Med Orb
Olfactory
Rolandic Oper
Front Inf Tri
Front Mid Orb
Front Sup Orb
Precentral
Tempr Pol Mid
Tempr Pol Sup
Heschl
Pallidum
Caudate
Precuneus
SupraMarginal
Parietal Sup
Fusiform
Occipital Mid
Lingual
Calcarine
ParaHippocamp
Cingulum Post
Insula
Rectus
Front Sup Med
Supp Motor Ar
Front Inf Orb
Front Inf Ope
Front Mid
Frontal Sup
-1
0
1
Mean activity
R
L
R
L
-1
-0.5
0
0.5
1
Mean activity
(normalised within state)

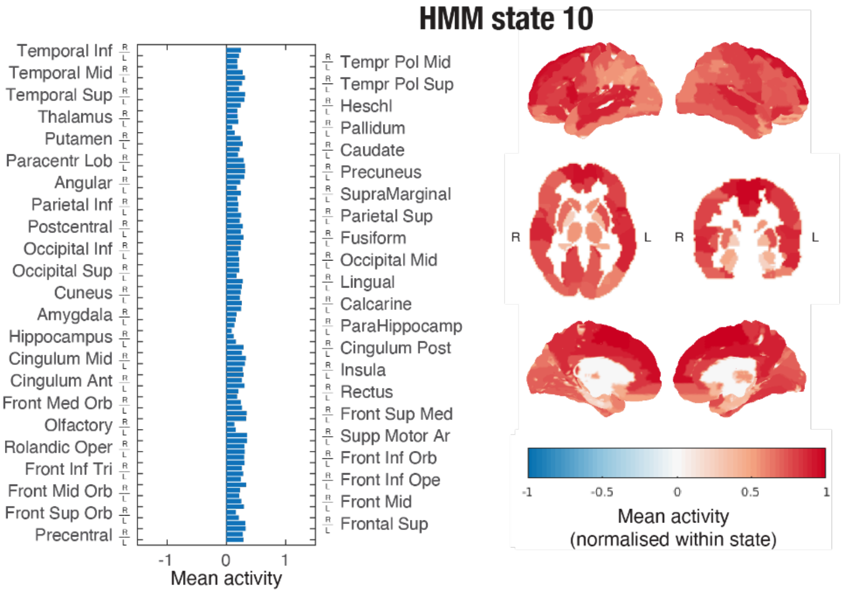


**Supplementary Figure S2** – The ten HMM states. For each HMM state is a bar plot of the mean activity in each of the AAL regions and a brain plot of the normalised activity for that given state.

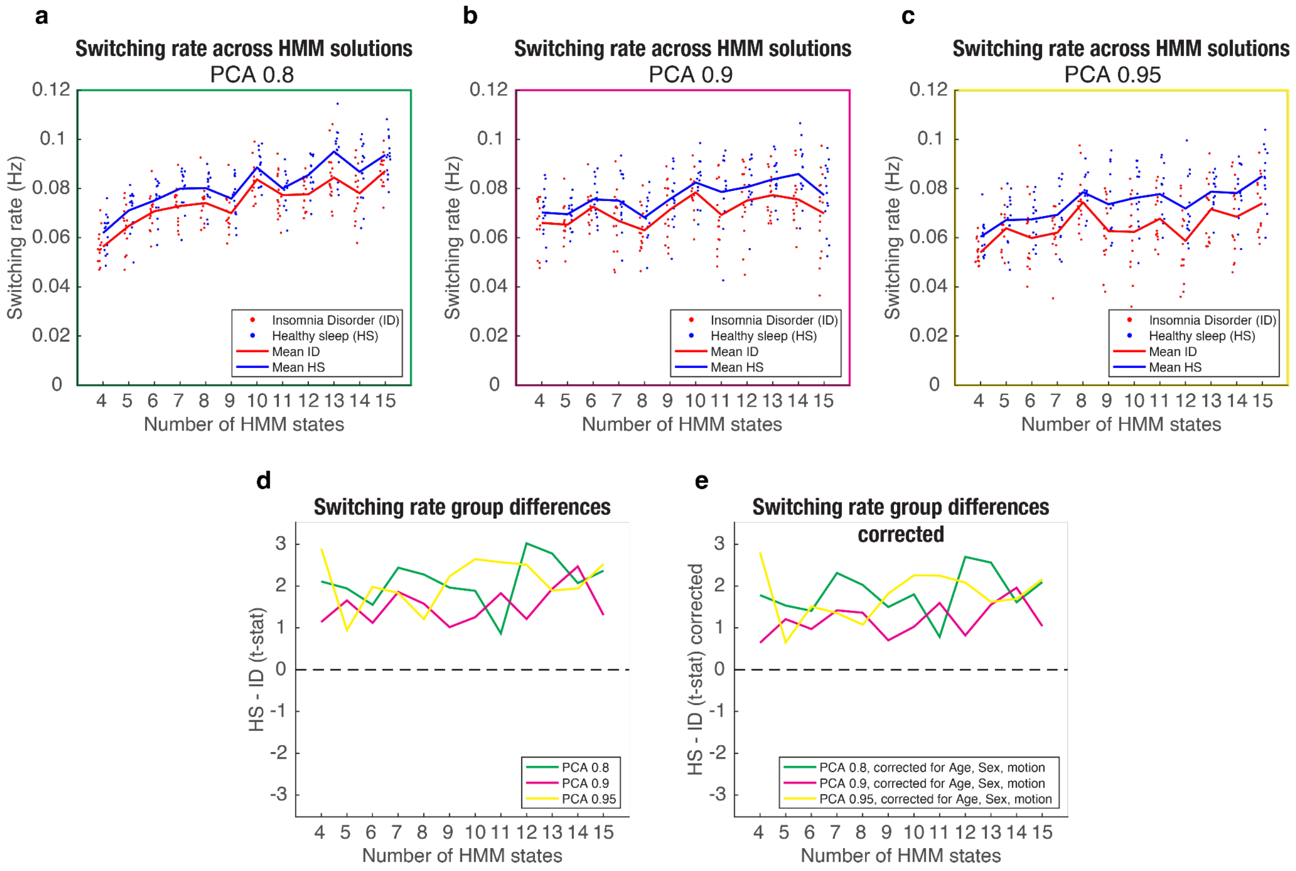


**Supplementary Figure S3** - Consistency in switching rate group difference between ID and HS across HMM solutions of varying PCA dimensionality and number of states. The between group differences in switching rate is similar across a range of number of states (4 to 15) for PCA dimensionality 0.8 (a), 0.9 (b) and 0.95 (c). Section C shows the direct comparison of the between group difference in switching rate for the three PCA dimensionalities, and section E shows the same comparison with a correction for age, sex and motion to validate that the group difference is not related to these three factors.

| | Model 1 | | | Model 2 | | | Model 3 | | |
|---|---|---|---|---|---|---|---|---|---|
| Linear model: | Switching ~ 1 + Sex + Age + ISI | | | Switching ~ 1 + Sex + Age + ISI + motionFMRI | | | Switching ~ 1 + Sex + Age + ISI + motionFMRI + Group:ISI | | |
| | *Estimate* | *(SE)* | *p-value* | *Estimate* | *(SE)* | *p-value* | *Estimate* | *(SE)* | *p-value* |
| Coefficients: | | | | | | | | | |
| (Intercept) | 0.081787 | (0.0094788) | 5.7606e-09 | 0.08686 | (0.0099794) | 6.8723e-09 | 0.087214 | (0.010255) | 1.4785e-08 |
| **Sex_Male** | -0.0074916 | (0.0056675) | 0.19818 | -0.00559 | (0.005724) | 0.33851 | -0.0050897 | (0.0061038) | 0.41294 |
| **Age** | -1.8998e-06 | (0.00019506) | 0.99231 | 2.6959e-06 | (0.00019142) | 0.98888 | 1.0618e-05 | (0.00019724) | 0.95753 |
| **ISI** | -0.00093304 | (0.00026369) | 0.001603 | -0.00077806 | (0.00028135) | 0.010755 | -0.00082314 | (0.00032888) | 0.019869 |
| **motionFMRI** | | | | -0.048904 | (0.034874) | 0.17363 | -0.048241 | (0.035642) | 0.18905 |
| **Group_HS:ISI** | | | | | | | -0.00036387 | (0.0012977) | 0.78169 |
| | Number of observations: 29, Error degrees of freedom: 25<br>Root Mean Squared Error: 0.0126<br>R-squared: 0.409, Adjusted R-Squared: 0.338<br>F-statistic vs. constant model: 5.77, p-value = 0.00386 | | | Number of observations: 29, Error degrees of freedom: 24<br>Root Mean Squared Error: 0.0123<br>R-squared: 0.454, Adjusted R-Squared: 0.363<br>F-statistic vs. constant model: 4.98, p-value = 0.00455 | | | Number of observations: 29, Error degrees of freedom: 23<br>Root Mean Squared Error: 0.0126<br>R-squared: 0.456, Adjusted R-Squared: 0.337<br>F-statistic vs. constant model: 3.85, p-value = 0.0111 | | |

**Table S1** - Results of the linear regression models. HS, healthy sleepers; ISI, Insomnia Severity Index.
